# Imaging seminiferous tubules – a 9.4T MRI mouse model

**DOI:** 10.1101/155762

**Authors:** M. Herigstad, S. Granados-Aparici, A. Pacey, M. Paley, S. Reynolds

## Abstract

Fertility problems affect many couples. Research into male fertility commonly uses mouse models due to their availability and similar spermatogenesis to humans. A common target is the seminiferous tubules, the site of spermatozoa production, typically studied using biopsies and histological analysis. High-field Magnetic Resonance (MR) may offer a non-invasive alternative to investigate testicular function in infertility models. Here, we assess seminiferous tubules structure in sacrificed mice to determine the usefulness of MR compared to histology. Twelve mice (11 aged 35-57 days, one >9 months) were sacrificed and MR imaged at 9.4T with a Rapid Acquisition with Relaxation Enhancement sequence. Testes were scanned in situ for all mice, and excised in a subset of mice (n=4). A second subset of mice (n=4) had their testes selected for histological analysis. Seminiferous tubule diameter was measured manually from MRI and histology images. Custom image analysis scripts were created for the automated segmentation of seminiferous tubules and calculation of tissue volumes. All ex vivo and in situ images of testes exhibited clear outlines of seminiferous tubules. Ratio of total testis volume to volume of seminiferous tubules did not differ significantly between ex vivo and in situ measurements, and were similar in mature and younger mice. Both total testis volume and seminiferous tubule volume were larger in the mature animal. While histological slices trended towards larger average seminiferous tubules diameter than MRI images, we found no significant differences between MRI and histological measurements. High-field MRI can be used in a mouse model to assess testicular structure in situ. All volumetric measurements compared favourably with histological data. In situ scans also clearly showed identifiable extra-testicular tissues, such as epididymis and prostate tissues. The potential to image tissues associated with sperm maturation as well as spermatogenesis emphasises how MR could be a useful technique in mouse models of fertility, however further work is required to optimize tissue segmentation and validate this method for use in longitudinal studies. This type of measurement could be extended to human fertility studies in the future.

## Introduction

The production of sperm during the reproductive period in male mammals requires several key steps, many of which are vulnerable to damage. Spermatogenesis is initiated in the epithelium of the seminiferous tubules, which are convoluted, tightly packed tubes that constitute most of the testis volume. Here, germ cells proliferate, go through meiosis and differentiation, before travelling via the rete testis into the epididymis, where they mature and gain the capacity for motility (1). Structure and function of the seminiferous tubules can be damaged by a range of medical conditions and environmental insults, including viral infections (2), diabetes (3), ischaemic insult (4, 5) and exposure to toxic compounds (6-8).

Mouse models are frequently used in male fertility research, due to their similarities with human spermatogenesis processes (1). Such models can be used to identify causes of defects or manipulate factors influencing key steps in spermatogenesis, which may not be practical nor ethical in human studies. Investigation of testis structure and function in mouse models of fertility typically requires dissection and histological analysis for visualisation and volume measurements of testes and their different sub-structures (e.g. seminiferous tubules, epididymis). These are invasive procedures necessitating the sacrifice of the animal, and are thus not compatible with longitudinal studies.

High-field magnetic resonance imaging (MRI) is non-invasive and has resolution capable of visualising the internal structures of mice testes. In this study, we investigate how MRI can be used to non-invasively assess testis size and structure in sacrificed mice, focusing on automated identification and segmentation of seminiferous tubules as a measure of testicular development. We seek to compare 9.4T MRI images of the mouse male reproductive system both in situ and ex vivo with histological assessments in the same testes. We will comment upon the usefulness of MRI in murine models of testicular health and disease and the potential for further development of this technique in human clinical investigations.

## Methods

### MRI protocol

Fig 1 illustrates the study protocol. Twelve sacrificed mice (11 aged 37-57 days, mean 41.6 ± 7.8, and one mature mouse (>9 months)) from strains with no associated testicular dysfunction were scanned in situ in a 9.4T MR scanner (Bruker BioSpin GmbH, Karlsruhe, Germany). The work was approved by the Project Applications and Amendments Committee (a sub-committee of the University of Sheffield Animal Welfare and Ethical Review Body). The animals were sacrificed using schedule 1 methods for reasons not related to this study. T1-weighted Fast Low Angle Shot (FLASH) sequences were obtained to identify the reproductive organs and plan for high-resolution imaging of testicular tissue. High-resolution imaging (in plane resolution 30-90μm) was performed with a Rapid Acquisition with Relaxation Enhancement (RARE) sequence (TR/TE 5000/28 ms, see Table 1 for full acquisition parameters). In a subset of mice (n=8), testes were excised. Out of these eight excised testes, four were re-imaged (one fresh and three after fixing in 10% buffered formalin). The other four were selected for histological analysis (see below). All animals and fresh ex vivo tissues were scanned immediately or within 24 hours of sacrifice (storage at -20^°^C), and fixed tissues were scanned within one week of sacrifice.

**Fig 1.**
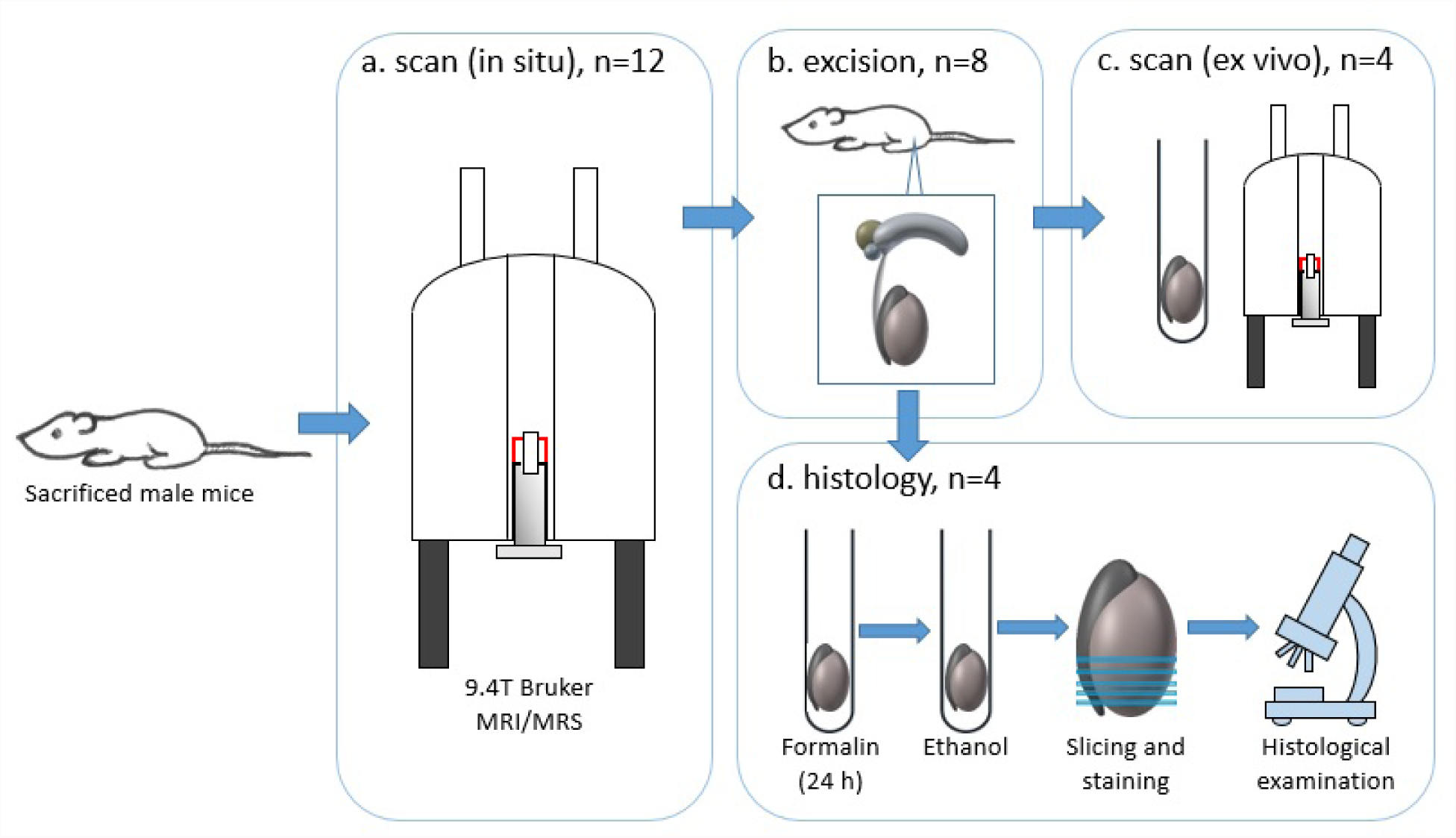
Schematic of protocol. Sacrificed male mice were first scanned in situ (a), followed by excision of testes in a subset of mice (b). Testes were then either scanned (c) or prepared for histological examination (d).

**Table 1:** Scan parameters

| Tissue | N | RARE<br>Factor | TE<br>(ms) | TR<br>(ms) | Slices/<br>Thickness<br>(mm) | NEX | Matrix | FOV (mm) | Resolution<br>(mm/px) | Scan<br>time<br>(min) |
| --- | --- | --- | --- | --- | --- | --- | --- | --- | --- | --- |
| In situ | N01 | 8 | 28 | 5000 | 27/0.25 | 8 | 256/256 | 14.1/17.5 | 0.055 | 42.40 |
|  | N02 | 4 | 28 | 5000 | 15/0.5 | 8 | 256/256 | 23/23 | 0.090 | 42.40 |
|  | N03 | 4 | 28 | 5000 | 17/0.5 | 8 | 300/300 | 25/25 | 0.083 | 50.00 |
|  | N04 | 4 | 28 | 5000 | 19/0.5 | 8 | 300/300 | 25/25 | 0.083 | 50.00 |
|  | N05 | 4 | 28 | 5000 | 29/0.5 | 50 | 522/533 | 23.5/24 | 0.045 | 554.10 |
|  | N06 | 4 | 28 | 5000 | 37/0.5 | 64 | 533/533 | 24/24 | 0.045 | 709.20 |
|  | N07 | 4 | 28 | 5000 | 33/0.5 | 32 | 556/489 | 25/22 | 0.045 | 325.2 |
|  | N08 | 4 | 28 | 5000 | 31/0.5 | 64 | 533/489 | 24/22 | 0.045 | 650.4 |
|  | N09 | 4 | 28 | 5000 | 21/0.5 | 8 | 500/390 | 22.5/17.5 | 0.045 | 64.4 |
|  | N10 | 4 | 28 | 5000 | 21/0.5 | 8 | 507/511 | 22.8/23 | 0.045 | 84.4 |
|  | N11 | 4 | 28 | 5000 | 23/1.0 | 8 | 533/523 | 24/23.5 | 0.045 | 86.4 |
|  | N12 | 4 | 28 | 5000 | 33/0.5 | 8 | 556/522 | 25/23.5 | 0.045 | 86.4 |
| Ex vivo | N01 | 8 | 28 | 5000 | 19/0.5 | 512 | 316/231 | 7.9/5.8 | 0.030 | 955.4 |
|  | N02 | 4 | 28 | 5000 | 19/0.5 | 180 | 356/240 | 10.4/7.3 | 0.030 | 900.0 |
|  | N03 | 4 | 28 | 5000 | 27/0.5 | 128 | 512/300 | 15.5/6 | 0.030 | 800.0 |
|  | N04 | 4 | 28 | 5000 | 19/1.0 | 128 | 433/300 | 13/9 | 0.030 | 800.0 |

N=subject number; RARE= Rapid Acquisition with Relaxation Enhancement; TE=time echo; TR=time repetition; NEX=number of averages; FOV=field of view.

### MR image analysis

Slices with clear seminiferous tubules were selected for each animal. Seminiferous tubule diameter was measured manually (single axis) using Paravision v5.1 (Bruker BioSpin). As tubular diameter is not uniform, an average of nine tubules across three slices (approximately centre-testis) were measured and the diameters averaged. Regions of interest (ROIs) were drawn around all slices judged to contain testicular tissue for one testis, chosen at random, from each animal. A custom MatLab (Mathworks, Natick, MA, USA) image analysis script was then used for segmentation of seminiferous tubules and calculation of tissue volumes. The script included three separate steps. First, full testis tissue volume was calculated based on the sum of all voxels in the ROI. Second, seminiferous tubules were segmented from testis tissue through thresholding (Otsu method (9)) to remove background noise. The segmentation was based on standard deviation (SD) of the mean signal within the ROI (low signal threshold: 1.5xSD, high signal threshold: 3xSD; thresholds were derived from preliminary analysis of excised tissue from mature animal). Third, seminiferous tissue volume was calculated based on the sum of the volumes of the non-zero voxels in the segmented images. The script was optimised using scans of excised tissues and subsequently implemented for the in situ scans.

### Histological preparation

Testes were excised and samples kept in 10% buffered formalin (NBS, Sigma, Irvine, UK) for 24 hours. The tissues were then dehydrated in an alcohol series to ensure complete water removal, treated with xylene and embedded in paraffin. Paraffin blocks were cut into 5μm sections and stained with hematoxylin and eosin (H&E, Surgipath Europe LTD, Peterborough, UK). Sections were imaged using 4X and 10X objectives (Olympus CKX41). Nine measurements of seminiferous tubule diameter were made across three sections (taken at approximately 50μm, 300μm and 550 μm from the testis edge) at 10X magnification (cross-sections of tubules) and the mean diameter calculated.

### Statistical analysis

Volumetric measurements were compared using a paired Student’s t-test, and the relationship between volumes and ages of the animals interrogated using linear correlation (MatLab). All values are quoted as mean ± SD.

## Results

### Images

Extra-testicular tissues were observed in situ using both FLASH and RARE imaging sequences and included the epididymis, prostate gland, bladder and penis (Fig 2). All animals yielded usable ex vivo and in situ images of testes, with clear outlines of seminiferous tubules (Fig 3). Histological analysis produced detailed images of testicular tissue with delineated seminiferous tubules (Fig 4). An example of identification of seminiferous tubules using automatic segmentation is shown in Fig 5.

**Fig 2.**
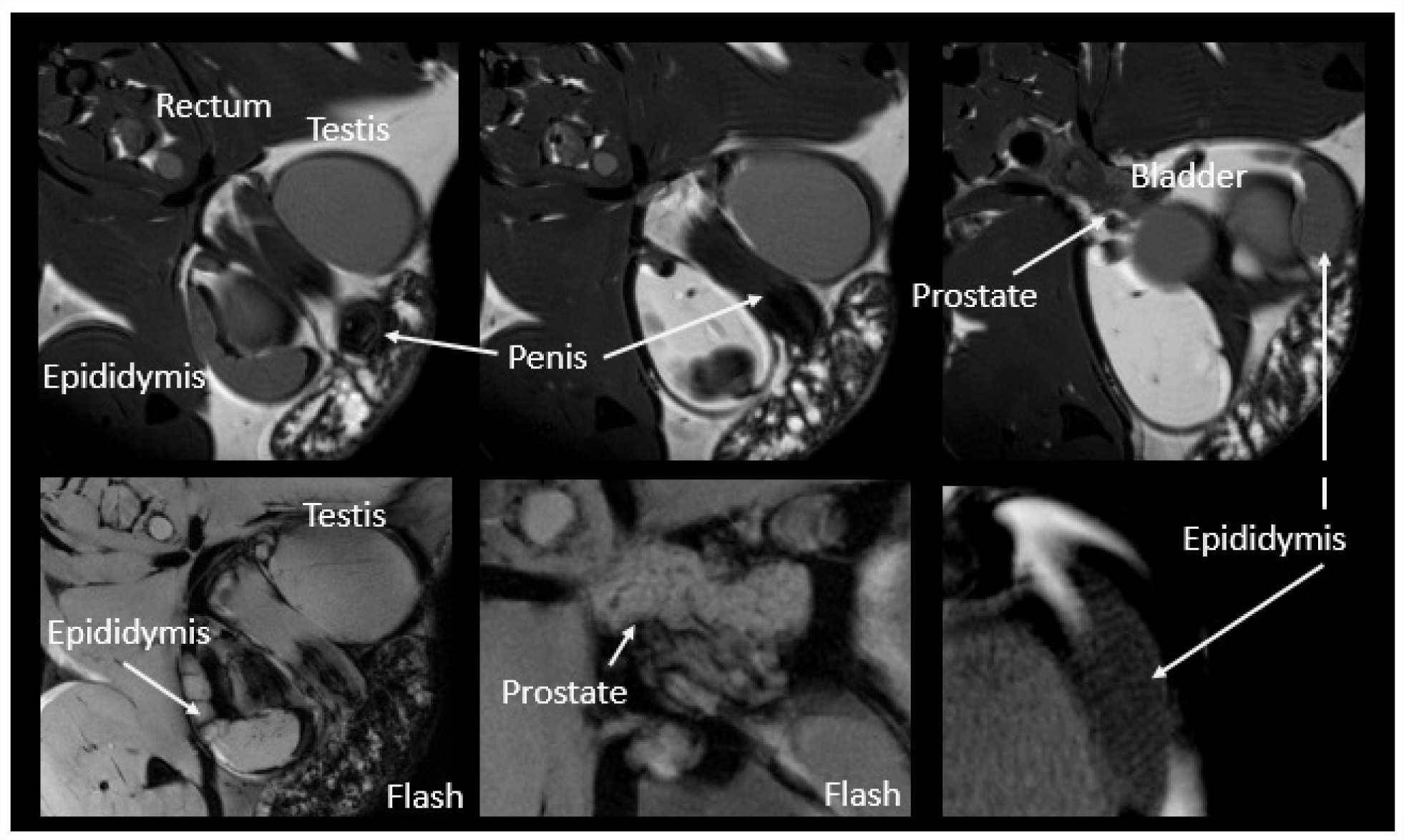
Testes and extra-testicular tissues obtained in situ. Images are from subject N5 and representative for the sample. Scan details are presented in Table 1. Images denoted ‘Flash’ were acquired using a T1-weighted Fast Low Angle Shot (FLASH) sequence (TE=8ms, TR=500ms, 32 averages, FOV=23.5x24mm, matrix=522/533).

**Fig 3.**
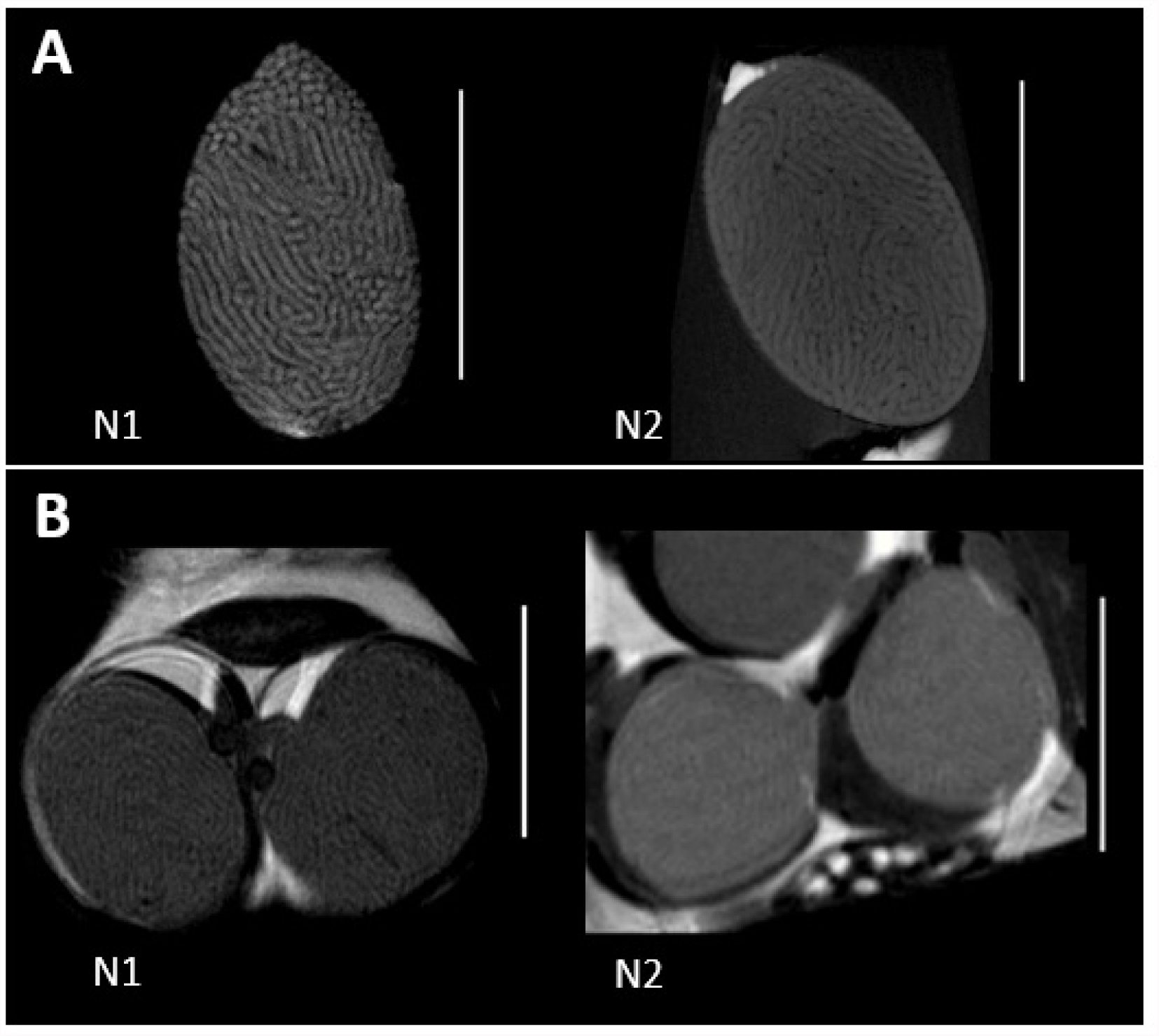
Structural MRI of mouse testes, ex vivo (A) and in situ (B). Slices approximately center-testis. Images are representative of the sample and include the mature reference individual (N1) and a younger animal (N2, age 35 days). Vertical white bars represent 5mm. Scan details are presented in Table 1.

**Fig 4.**
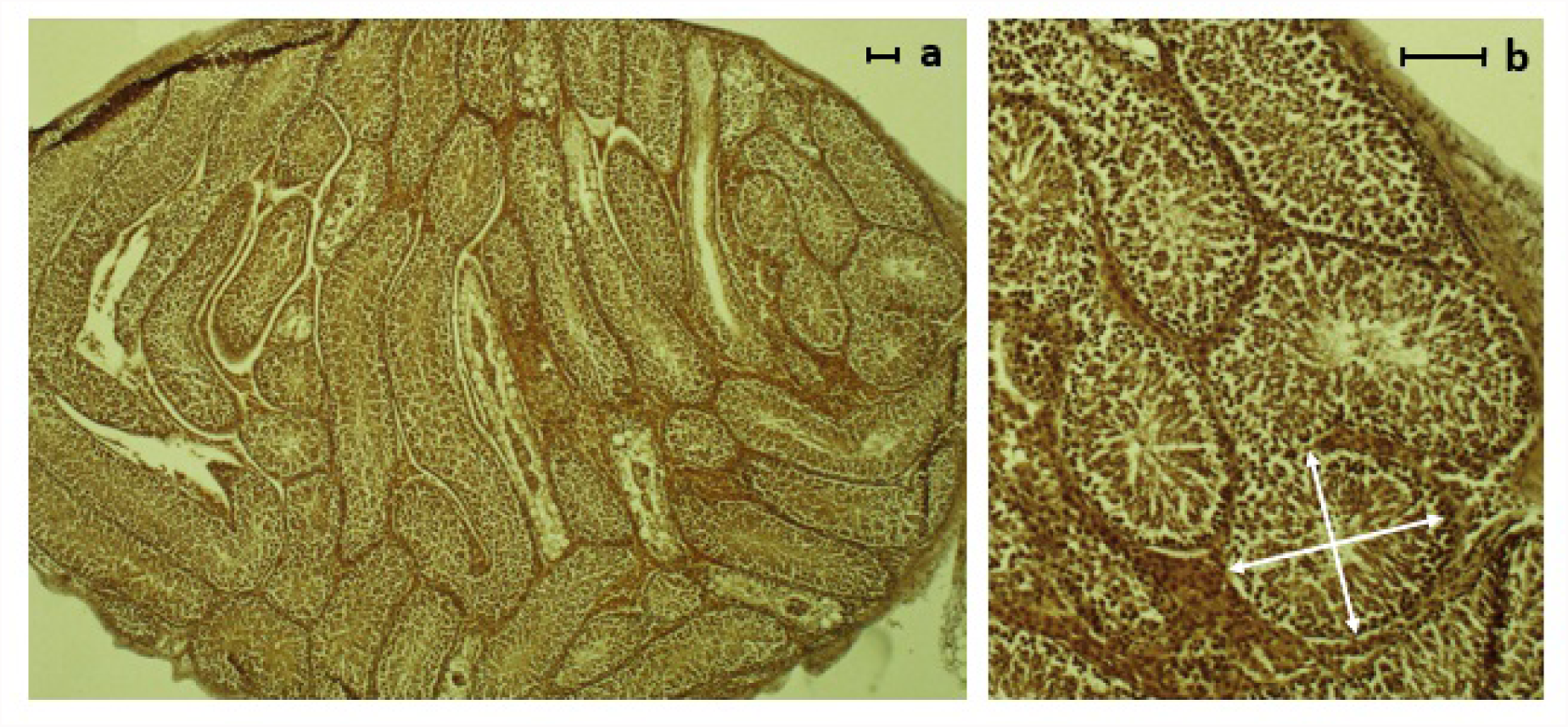
H&E staining of testes. Images are from subject N11 and representative for the sample. a=4x magnification, b=10x magnification. Scale bars (right corners) both 100μm. White arrows showing example of cross section used for measurements (10).

**Fig 5.**
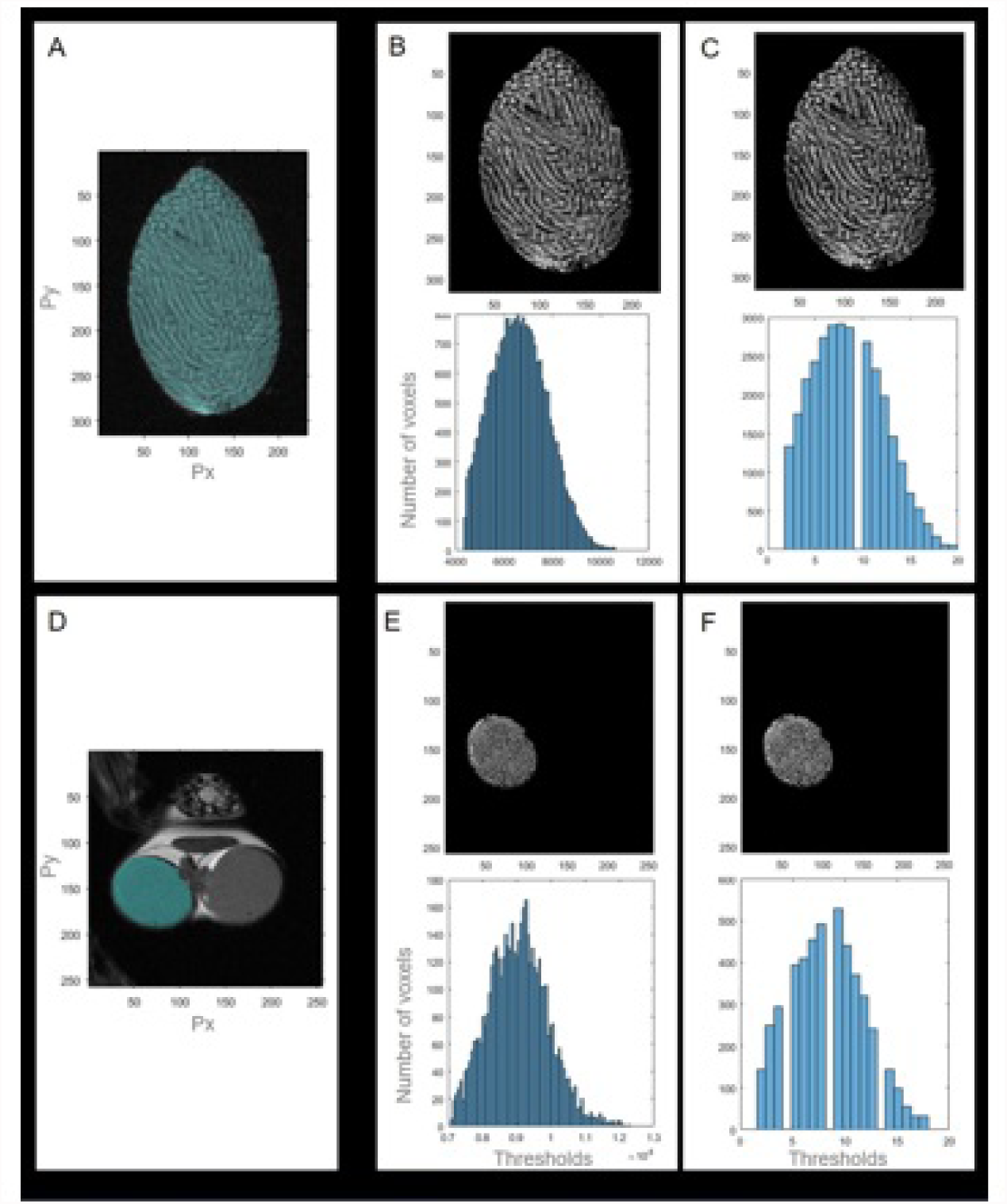
Example of identification of seminiferous tubules using segmentation in subject N1. Regions of interest were drawn around testis tissue (A and D). Images were thresholded (B and E) using the Otsu method in MatLab, with high and low thresholds assigned based on standard deviation of the mean signal (low 1.5xSD, high 3xSD). Seminiferous tubules and total tissue volume was determined based on the sum of voxels in the processed images. Histogram represents number of voxels at the threshold level used. A-C, ex vivo; D-F, in situ. C and F, stratified images.

### Calculated volumes

Volumes were calculated for in situ (n=12) and ex vivo scans (n=4). In situ scans showed a total testis volume of 63.2 ± 7.6μl (mean ± SD), a seminiferous tubules volume of 57.0 ± 6.9μl, and a seminiferous tubule volume to total testis volume ratio of 0.90 ± 0.01.

Comparing the scans in mice selected for ex vivo MRI (n=4, one fresh and three fixed in 10% buffered formalin), we observed no significant differences between in situ and ex vivo measurements. In these animals, the total testis volume was 69.9 ± 5.5μl in situ and 69.9 ± 2.2μl ex vivo (p=0.97), the seminiferous tubules volume was 63.1 ± 5.1μl in situ and 63.2 ± 1.7μl ex vivo (p=0.99). One animal (N3) showed lower volume (but no change in volume ratio) ex vivo due to compressed tissues in the sample tube and was excluded from the comparison. Ratio calculations showed no difference in situ versus ex vivo (both 0.90 ± 0.01, p=0.94).

### Seminiferous tubules diameter

Average seminiferous tubule diameter measured from in situ RARE images was 164 ± 16μm (range 141-198, n=12). Comparing measurements of seminiferous tubule diameter in the mice selected for histological analysis (n=4), we observed no significant differences between averages obtained using histology and using MRI. MRI analysis in this subset showed average tubule diameter of 156 ± 7μm (range 147-163, in situ images). Histological analysis in the same animals showed larger diameters (169 ± 7μm, range 160-175), but this difference did not reach statistical significance (p=0.11).

### Age effects

Total testis and duct volume was larger in the mature reference animal compared to the mean of the younger individuals by 19.7% and 19.8% respectively (no statistical comparisons). Calculated total testis volume to seminiferous tubules ratio was the same in the mature animal as younger mice, but the mean width of the seminiferous tubules were larger size in the mature animal (12.2%, no statistical comparisons). There were no significant correlations between age and testes or seminiferous volumetric measurements within the main sample (aged 37-57 days).

## Discussion

### Key findings

Mouse seminiferous tubules structure and extra-testicular tissues may be visualised in situ using 9.4T MRI. MR images may be analysed and volumetric information quantified using automated image analysis scripts.

### Quantification of images

The automated image analysis yielded data compatible with previous histological findings in similar animal populations (10-12). We observed that total testis size was greater in the mature individual compared to younger mice (10), but there was predictably no significant correlation with age in the main sample, in which the majority of the mice were of a similar age (35-38 days old, N=7). The volume of seminiferous tubules was matched to testicular volume, as the total testicular volume to tubules ratio was similar between individuals irrespective of testes size. This suggests no change in testicular architecture within the age group (10). In the subset of mice selected for ex vivo imaging, the calculated volumes did not differ between ex vivo and in situ scans. Our quantification method thus yields repeatable and reliable volume estimates. Furthermore, as in situ scans were collected over a shorter period and with fewer averages than ex vivo scans (Table 1), this indicates that our analysis method remains robust across different levels of contrast.

We obtained in situ scans with in-plane resolution of 45μm (slice thickness 0.5mm), which allowed us to observe the tubules (typically 150-200μm in diameter). Manual estimates of tubule diameter were not significantly different to, albeit trending towards lower values than, those observed using histology. Lower resolution and partial volume effects in the MR images may account in part for this trend. Additionally, as thresholding was used to segment seminiferous tubules to obtain a clearer view of the testicular composition, this could cause edges of tubules to be consistently underestimated as a single voxel width was used to delineate adjacent tubules. Interpolating the images may improve the accuracy of diameter estimation. While histology remains the gold standard for measuring absolute tissue morphometry in small structures, MRI might be more accurate in larger structures as it circumvents sources of tissue distortion (e.g. in murine brain imaging (13)).

MRI has a wider Field of View capability than most invasive procedures (e.g. histological analysis), permitting the 3D exploration of whole structures with excellent soft tissue contrast. Comparatively, histological analysis is limited to selected, small slices. MRI by contrast can capture full 3D images of structures that would otherwise be too large for standard histological analysis. Here, we reliably obtained clear structural images of urogenital components, including the epididymis, penis, prostate and bladder (Fig 3). The epididymis is of particular interest for fertility research, as sperm maturation and development of motility potential occurs in this tissue (14). This unique temporal and spatial capability makes MRI a valuable tool, and it has already been used in a range of murine models of human disease (13, 15, 16) and physiology (17-19). Yet it remains underutilised in fertility research.

At 9.4T it is possible to obtain greater resolution and contrast than those obtained here, although this comes at the cost of extended scan time. Fig 3 shows ex vivo and in situ images of testes from two animals, illustrating the added contrast obtained in longer scans (ex vivo) compared to shorter scans (in situ). Despite this difference, our automated volume calculations were highly similar for in situ and ex vivo scans. Furthermore, T2 weighted fast spin-echo images with resolution of 30μm has been achieved in 60 minutes for murine in vivo experiments without contrast agents (20). In rat testes, in vivo scans at lower field strengths (4.7T) show tubular structure in T2 weighted images with scan times of 68 minutes or less (5). Even lower field strengths (1.5T) have successfully been used in rats to detect hypoperfusion following experimental testicular torsion using scan sequences lasting less than 10 minutes (5, 21). This, combined with our high-resolution structural images in situ, strongly suggests that high field strength MRI is both feasible and useful when investigating murine testes in vivo.

### MRI in murine models of fertility

Translation from animal research to human medicine is greatly aided by longitudinal experimental designs. Such protocols add a temporal component to pre-clinical research that is generally considered central to fully characterise physiological processes. MRI is uniquely positioned to conduct this type of research as its non-invasive nature permits the study of progressions of disease or development within-subject, resulting in increased experimental power and the potential for using (and sacrificing) fewer animals. This is in line with the principle of the 3Rs in animal research: Replacement, Reduction and Refinement.

The potential of MR extends beyond structural imaging, as live mice offer a range of additional functional data. Diffusion imaging or functional imaging may provide information on fluid movement and blood perfusion. Regional ischaemia, heightened perfusion, bloating vessels, and blocked or damaged seminiferous tubules can all be imaged at high resolution. Contrast agents may be employed to further optimise contrast and track vascularisation. Magnetic resonance spectroscopy (MRS) information can also be obtained. MRS is a highly powerful tool for metabolic research that may allow us to probe spatially discreet metabolism. For example, MRS might be used to investigate testicular tumour metabolism in vivo, or to track spermatogenesis variation over time within seminiferous tubules.

### Clinical potential

MRI/MRS may also be valuable clinical tools. For example, MRI can be used in humans to visualise intratesticular lesions (22-24), testicular tumours (25), torsion (21, 26), varicoceles (27, 28) and testicle location (29) at common clinical MR field strengths. Reduction in testicular size, epididymis diameter or seminiferous tubule diameter (30) may be markers of disrupted spermatogenesis, and the size of either could thus potentially be used as an indicator of pathology. This is supported by evidence from MR images in rats with chronic spermatogenic impairment (5). MRI assessments may also be used to guide testicular biopsies (31), reducing the number of invasive events and thus lessen the chance of surgical complications and infertility (32). Furthermore, recent findings suggest that MRS may be used to assess metabolite concentration in the testes in both humans (33) and rats (34), which could offer information on spermatogenesis (33), possibly through assessment of phosphocholine concentration as suggested by ^1^H magnetic resonance spectroscopy of snap frozen testicular tissue from men with azoospermia (35). MR could thus be a useful technique in the clinic as well as a valuable tool for experimental models of fertility.

### Limitations

The present study was conducted in deceased animals, and tissues were thus ischaemic at the point of scanning. Some tissue degradation and blood pooling may also have occurred despite scanning commencing immediately after sacrifice. Both ischaemia and tissue degradation would contribute to reduced image quality.

## Conclusions

Damage to the testes may significantly impair fertility, and the investigation of testis structure is central to research in male fertility. Short breeding cycles along with comparable spermatogenesis to humans and a range of relevant breeds make the mouse model highly useful. High-field MRI is a useful tool for mouse models of testicular morphology. Testicular composition can be visualised, and volumetric analysis of MR images is predictable and reproducible for a murine model exhibiting no testicular dysmorphy. The potential to image tissues associated with sperm maturation as well as spermatogenesis emphasizes how MR could be a useful technique in mouse models of fertility.

## Acknowledgements

We would like to thank the staff at the Biological Services Unit for their generous help and for supplying the animals for this study. The study was funded by the MRC (Grant number MR/M010473-1).

